# Conserved Structural Motifs in the Hammerhead Ribozyme of a Chloroplast Viroid Mimic tRNA Anticodon Structure to Hijack tRNA Ligase for Viroid Circularization

**DOI:** 10.1101/2022.01.19.477025

**Authors:** Beltrán Ortolá, José-Antonio Daròs

## Abstract

Viroids belonging to the family *Avsunviroidae* contain hammerhead ribozymes that process to unit length the oligomeric RNAs of both polarities generated during the rolling-circle replication that occurs in chloroplasts of host plants. Linear products, with 5’-hydroxyl and 2’,3’-phosphodiester termini, are then recognized and circularized by the host chloroplastic isoform of the tRNA ligase. Here we analyze the circularization process of eggplant latent viroid (ELVd), an asymptomatic viroid that infects eggplants (*Solanum melongena* L.), using an *Escherichia coli* co-expression system in which longer-than-unit linear ELVd (+) precursors are expressed along with the eggplant chloroplastic tRNA ligase. The RNA precursor contains two copies of the hammerhead ribozyme and yields the appropriate termini for the tRNA ligase-mediated ligation in bacteria. We have determined that the ligation efficiency is highly dependent on the presence of ribozyme sequences in the ligatable termini, since the circularization of a series of viroid variants in which the ligation position was rearranged increased substantially in the presence of these sequences. Further *in silico* analysis showed sequence and structure similarity between the hammerhead ribozyme catalytic pocket and the anticodon loop of tRNAs, both of which harbor a characteristic U-turn of the phosphodiester backbone. Directed mutagenesis in the ribozyme domain supports the role of this U-turn loop in the ligation process. We propose that, in addition to its self-cleavage function, the viroid ribozymes have evolved to mimic the structure of the tRNA anticodon loop to recruit host tRNA ligase for the circularization of the monomeric linear replication intermediates.

**IMPORTANCE:** Viroids are a very particular class of infectious agents because they only consist of a small RNA that, to our current knowledge, does not encode for proteins. Consequently, viroids parasite host factors and structures to mediate all processes in the infectious cycle. How these small infectious RNAs are able to hijack host resources is currently a mystery. In this work, we shed some light on the functionality of hammerhead ribozymes during replication of viroids that belong to the family *Avsunviroidae*, which replicate in the chloroplasts. Our findings suggest that, in addition to mediate self-cleavage of replication intermediates, hammerhead ribozymes also recruit tRNA ligase for monomer circularization, likely mimicking a common host tRNA structural motif.

---

Viroids, consisting of small, circular, single-stranded, non-coding RNA molecules ranging from 246 to 434 nucleotides (nt), are the smallest known infectious agents (Adkar-Purushothama and Perreault, 2020; Matsushita et al., 2018; Navarro et al., 2021; Wang, 2021). Viroids rely on their structural RNA elements to hijack the appropriate cellular machinery to successfully infect certain higher plants, where they complete their replicative cycles. Using mechanisms that are not yet fully understood, they are able to move between cellular compartments and through plasmodesmata and phloem to establish systemic infections, escaping the host defensive response and occasionally causing economically important diseases.

Since most, if not all, processes in viroid’s replication cycle are based on interactions between viroid RNA structures and host elements, the presence or absence of certain conserved structural domains and motifs correlates with important physiological differences among the different species; these differences have been used in taxonomy (Di Serio et al., 2014). The most common and first-described viroids have a functional central conserved region (CCR) in their molecules and are grouped together within the family *Pospiviroidae*. These viroids replicate through the asymmetric variant of a rolling circle mechanism (Branch et al., 1988; Branch and Robertson, 1984). They are imported into the nucleus (Diener, 1971; Spiesmacher et al., 1983), where they are transcribed by the host DNA-dependent RNA polymerase II to produce lineal concatemers of complementary polarity (Mühlbach and Sänger, 1979). In viroids, + polarity is arbitrarily attributed to the most abundant circular RNA. Viroid concatemers of - polarity enter directly into a second round of transcription to form a complementary concatemer with + polarity. Host factors, such as a particular splicing form of transcription factor IIIA (TFIIIA-7ZF) and ribosomal protein L5, acting as a splicing regulator, are also involved in the replication process (Jiang et al., 2018). Finally, CCR elements are recognized and processed by a host RNase III and DNA ligase 1 to produce the viroid monomeric circular progeny (Gas et al., 2008, 2007; M. Á. Nohales et al., 2012).

Only a few viroids comprise the family *Avsunviroidae* (Di Serio et al., 2018; Flores et al., 2000). They lack a CCR but do contain functional hammerhead ribozymes in the strands of both polarities. The members of this family replicate through the symmetric variant of the rolling-circle mechanism (Branch and Robertson, 1984; Daròs et al., 1994), after they are imported into the chloroplast (Bonfiglioli et al., 1994), although they may still have a nuclear phase (Gómez and Pallás, 2012). In this family, the viroid molecules with + polarity serve as templates in a rolling-circle transcription to produce lineal concatemers with - polarity via a chloroplastic nuclear-encoded polymerase (Navarro et al., 2000). The action of the hammerhead ribozymes present in these concatemers produces monomeric intermediates with 5’-hydroxyl and 2’,3’-cyclic phosphodiester termini (Daròs and Flores, 2002). These termini are involved in the formation of an intramolecular 5’,3’-phosphodiester linkage to generate circular molecules with - polarity that serve as templates in a second rolling-circle transcription, symmetric to the first, that produces concatemers with + polarity, which also self-cleave through hammerhead ribozymes and then circularize to generate the circular progeny with + polarity.

Initially, ribozymes themselves were believed to circularize viroids; however, given the low efficiency of the backwards reaction and the formation of 2’-5’ linkages, the participation of a chloroplastic enzyme was suggested (Martínez et al., 2009). However, for several years, the presence in chloroplasts of enzymes with such properties was unknown. The demonstration of enzyme intervention in the circularization process within the family *Avsunviroidae* came from studies with the eggplant latent viroid (ELVd), an asymptomatic infectious agent of eggplants (*Solanum melongena* L.) that is the only species of the genus *Elaviroid* (Daròs, 2016; Fadda et al., 2003). If ELVd is expressed as a dimeric transcript in chloroplasts of the unicellular green alga *Chlamydomonas reinhardtii* (phylum Chlorophyta), it is processed into monomers and recognized by an enzyme that efficiently circularize the viroid (Martínez et al., 2009; Molina-Serrano et al., 2007). Moreover, with the unexpected discovery that tRNA splicing machinery is targeted to multiple cellular compartments in plants, including chloroplasts (Englert et al., 2007), the chloroplastic isoform of the eggplant tRNA ligase was identified as the host enzyme involved in the circularization of ELVd (and probably of all viroids in the family *Avsunviroidae*) (M.-A. Nohales et al., 2012). Furthermore, these studies highlighted the important role that a quasi-double-stranded structure present in the central part of the ELVd molecule plays in the ligation process (Martínez et al., 2009), along with other domains that are dispensable for ligation (Daròs et al., 2018). Because this central region results from the hybridization of the ribozyme domains of both polarities, it has been proposed that ribozyme sequences and/or structures play critical roles in the ligation of the monomeric linear replication intermediates, in addition to self-cleavage (Cordero et al., 2018). Experimental support to this hypothesis was accomplished by taking advantage of an *Escherichia coli* experimental system that allows for the accumulation of circular ELVd molecules in the bacteria after the co-expression of ELVd longer-than-unit transcripts and the chloroplastic isoform of the eggplant tRNA ligase (Cordero et al., 2018; Daròs et al., 2018). In this experimental system, the viroid expression cassette contains two copies of the hammerhead ribozyme cDNA surrounding the rest of the viroid sequence, allowing it to mimic natural viroid processing, as it generates a longer-than-unit viroid transcript in which the activity of the flanking ribozymes produces the appropriate termini for circularization by the co-expressed eggplant tRNA ligase.

Here, we use this *E. coli* co-expression system to gain insight into the ELVd sequence and structure requirements for recruiting the eggplant chloroplastic tRNA ligase to accomplish viroid circularization. By producing in *E. coli* a series of ELVd monomeric linear intermediates–with the ligatable 5’-hydroxyl and 2’,3’-cyclic phosphodiester termini opened at different positions in the molecule–we found that the efficiency of eggplant tRNA ligase-mediated ligation correlates with the presence of ribozyme sequences at the ligation site. In addition, our analysis of the hammerhead ribozyme domain reveals similarity between the motif that houses some of the conserved catalytic sequences and the seven nucleotides of the tRNA anticodon loop, including a uridine sharp turn that is functionally relevant in both domains. Our analysis supports that efficient ligation depends on the presence of these conserved sequences in the motif. Here, we propose a model in which viroid hammerhead ribozymes have evolutionarily acquired the double function of self-cleavage and of recruiting the host tRNA ligase by mimicking the host tRNA ligation site.

## RESULTS

### Circularization of ELVd rearranged forms by the eggplant tRNA ligase

To better understand the requirements for ELVd circularization by the chloroplastic isoform of eggplant tRNA ligase, we co-expressed this enzyme in *E. coli* along with different forms of the viroid monomeric linear replication intermediate opened at different positions in the RNA molecule. The opening sites are depicted in Fig. 1A based on an ELVd secondary structure previously determined experimentally (López-Carrasco et al., 2016). All ELVd forms contained the 5’-hydroxyl and 2’,3’-cyclic phosphodiester termini that are required for circularization by the tRNA ligase, as they resulted from processing of longer-than-unit precursors by flanking engineered ribozymes (Fig. 1B and Supplementary Dataset 1). We analyzed six monomeric linear ELVd forms whose circularization by eggplant tRNA ligase had previously been studied *in vitro* (M.-A. Nohales et al., 2012). Their opening sites were distributed throughout the entire viroid molecule within regions with different secondary structures and with different terminal nucleotides (Fig. 1A). To avoid internal self-cleavage of these monomeric linear ELVd forms by their endogenous hammerhead ribozyme, the strictly conserved CUGA box was mutated to UUGG in all of them (Fig. 1B and Supplementary Dataset 1).

**FIG 1.**
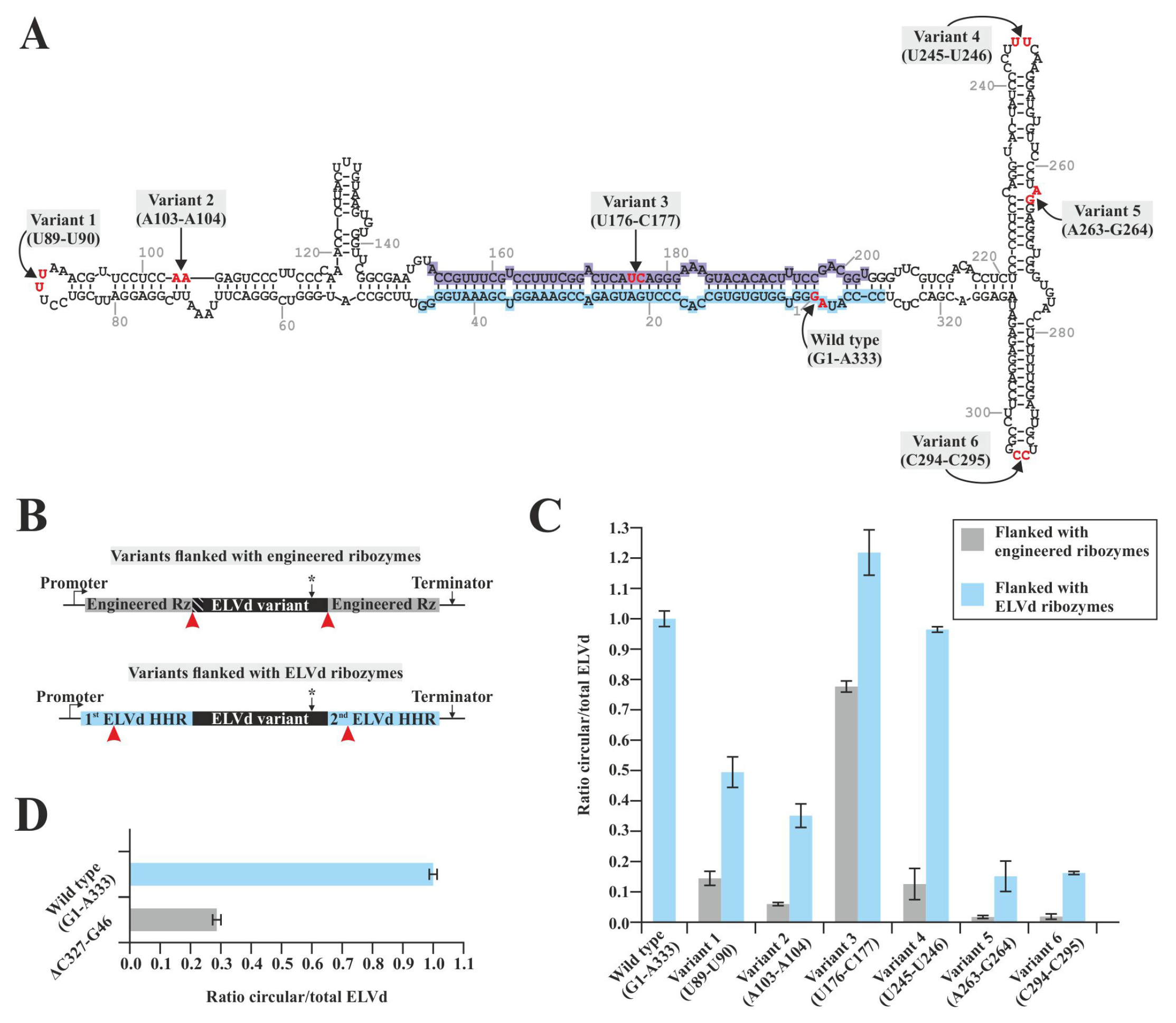
Eggplant tRNA ligase-mediated circularization of different monomeric linear ELVd (+) RNAs opened at different sites. (**A**) Opening sites of wild-type ELVd (G1-A333) and Variants 1 to 6 (U89-U90, A103-A104, U176-C177, U245-U246, A263-G264, and C294-C295, respectively) mapped onto the structure of the monomeric (+) circular ELVd. The domains of the hammerhead ribozymes of both polarities are highlighted in ice blue (+) and pastel blue (-). (**B**) For each variant, two constructs were generated, using either engineered ribozymes or the ELVd + ribozyme. Cleavage sites are indicated by red arrowheads. The mutation in the endogenous ribozyme CUGA is represented with an asterisk. Schematic representation is not at scale. The RNA precursors were co-expressed in *E. coli* along with the tRNA ligase; the total RNA from the bacteria was separated by 2D-PAGE and transferred to a membrane for northern blot hybridization with an ELVd (-) probe. (**C**) Histogram showing the normalized accumulation rate of monomeric circular versus monomeric total (circular plus linear) ELVd (+) of RNA variants flanked by engineered ribozymes (gray bars) or ELVd + ribozymes (blue bars). (**D**) Analysis of the circularization of an ELVd deletion mutant lacking the ribozyme sequence (ΔC327-G46) and including engineered ribozymes. (**C** and **D**) Error bars represent the standard deviations in three independent *E. coli* clones.

For each ELVd form, three independent co-transformed *E. coli* clones were grown in liquid cultures, and total RNA was extracted after 24 h. Viroid monomeric circular and linear molecules were separated using two-dimensional (2D) polyacrylamide gel electrophoresis (PAGE) and quantified via northern blot hybridization with a complementary ^32^P-labelled RNA probe (Supplementary Figure S1). The circularization rate (monomeric circular forms divided by total monomeric forms [linear plus circular]) was calculated from hybridization signals. The results show that circularization was substantially reduced in all rearranged ELVd forms (Fig. 1C, gray bars), as compared to wild-type ELVd opened at the genuine position (A333-G1). Most rearranged forms circularized at a rate of approximately 15% or less with respect to the wild type (Fig. 1C); the exception was Variant 3 (U176-C177), which was opened at the upper strand of the central quasi-double-stranded structure (Fig. 1A), which circularized about 78% of the wild type (Fig. 1C). The strong reduction in circularization of most reorganized variants precludes the possibility that only the terminal 5’-hydroxyl and 2’,3’-cyclic phosphodiester groups are required for efficient tRNA ligase-mediated ligation of linear ELVd. Interestingly, the only form that was substantially circularized (Variant 3) is opened at the center of the ELVd molecule, relatively close to the genuine circularization site. This opening site maps in a domain that corresponds to the hammerhead ribozyme of complementary (-) polarity, opposite to that of the + strand in the viroid secondary structure (Fig. 1A). Overall, these results suggest that the sequences or structures of the hammerhead ribozyme in the vicinity of the ligation site may favor the recognition of linear ELVd intermediates by the eggplant tRNA ligase.

### Effect of the hammerhead ribozyme on tRNA ligase-mediated ELVd circularization

To test this hypothesis, we analyzed whether the presence of sequences corresponding to the ELVd (+) hammerhead ribozyme at both termini of the different rearranged monomeric linear viroid forms affects circularization by the tRNA ligase. To this end, we built a new set of plasmids to express in *E. coli* the same ELVd rearranged forms, but now flanked on both sides by the whole domain of the viroid ribozyme with + polarity (Fig. 1B and Supplementary Dataset 1). Notably, although this approach returns the native sequence to the ligation sites, it also results in the insertion of 53 nt (the full ribozyme domain) in different positions within the viroid molecule. Again, we grew liquid cultures from three independent *E. coli* clones co-transformed to co-express the eggplant tRNA ligase and the different ELVd forms; viroid RNA was analyzed at 24 h. Remarkably, the ratio of circularization increased significantly in all rearranged viroid forms when flanked by the hammerhead ribozyme sequences (Fig. 1C, blue bars). Interestingly, the ratio of circularization of Variant 4 (U245-U246) was now close to that of the wild type, which was even surpassed in the case of Variant 3 (U176-C177) (Fig. 1C, compare blue bars with that corresponding to wild-type). We also analyzed the effect of deleting the hammerhead ribozyme in the wild-type monomeric linear intermediate (Supplementary Dataset 1). Circularization was drastically reduced to approximately 28% when replacing flanking native hammerhead ribozymes from the wild-type viroid by engineered ribozymes (Fig. 1D).

To further analyze the effect of hammerhead ribozyme on tRNA ligase-mediated circularization of ELVd, we focused on Variant 2 (A103-A104), which initially displayed a moderately low rate of ligation (~6%) that increased to roughly 35% when the viroid ribozyme halves were added to the terminal ends (Fig. 1C). Based on the version of Variant 2 flanked at both sides by the viroid ribozymes, we built a new set of plasmids to express monomeric linear forms (2A to D) in *E. coli*. In these forms, the added ribozymes were extended by 20 nt at the 3’ end (2A), the 5’ end (2B), or both ends (2C); alternatively, the sequence complementary to the viroid ribozyme with - polarity was inserted between positions U72 and U73 (2D) to mimic the secondary structure of wild-type ELVd (Fig. 2A and Supplementary Dataset 1). Expression of these viroid forms in *E. coli* along the eggplant tRNA ligase generated increased rates of ligation compared to Variant 2, in which strict viroid ribozyme was added (Fig. 2B). The 20-nt 3’ extension of the ribozyme brought circularization rates close to those of wild-type ELVd, while the 20-nt 5’ extension increased the circularization rate from 35% to approximately 61%. Both extensions together displayed an additive effect, with a rate of ligation that surpassed that of the wild type. Finally, the insertion of the - ribozyme domain in the opposite strand increased the rate to approximately 80% (Fig. 2B). Together, these results indicate that the re-creation of the genuine ligation site in the ELVd molecule dramatically improves tRNA ligase-mediated ligation.

**FIG 2.**
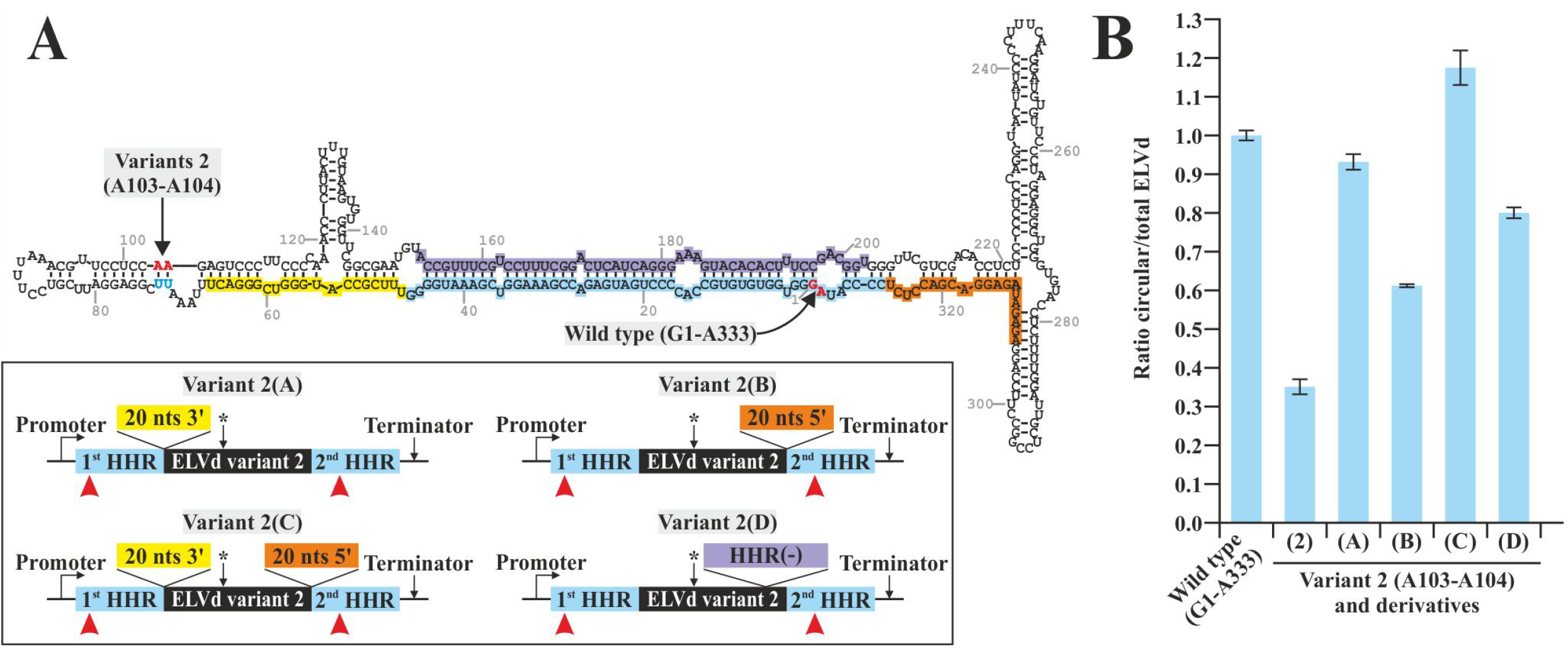
Analysis of the role of sequences surrounding the hammerhead ribozyme on the monomeric linear ELVd (+) intermediate ligation by the eggplant tRNA ligase in *E. coli*. (**A**) New set of mutants based on modifications in Variant 2 (opened at C294-C295), containing ELVd hammerhead ribozyme (+) halves in both termini. A schematic representation (not at scale) of Variants 2A, B, C, and D is shown in the box. Variants 2A, B, and C contained a 20-nt extension of their 3’ end, 5’ end, or both 3’ and 5’ ends, respectively. Variant 2D contained the insertion of the domain corresponding to the ribozyme with - polarity inserted between positions U72 and U73 (in blue). The sequences of hammerhead ribozymes of both polarities are ice blue (+) and pastel blue (-). Cleavage sites of the hammerhead ribozymes are indicated by red arrowheads. The mutation in the endogenous CUGA sequence is represented with an asterisk. (**B**) Circularization rates were analyzed by northern blot hybridization of 2D separated RNA from *E. coli* clones, in which the ELVd variants were co-expressed with eggplant tRNA ligase. Histogram showing the accumulation rate of monomeric circular versus monomeric total (circular and linear) ELVd (+) RNA in the indicated variants. Error bars represent the standard deviations in three independent *E. coli* clones.

### Effect of mutating the viroid hammerhead ribozyme on tRNA ligase-mediated ELVd circularization

Because circularization of ELVd forms increases when ligation sites reside in (or are in the vicinity of) the sequences that conform the hammerhead ribozyme, we hypothesized that, in addition to self-cleavage, the sequences in this domain may also be involved in recognition by the eggplant tRNA ligase, perhaps by mimicking *bonafide* tRNA ligase substrates, such as the tRNA halves. To further investigate this hypothesis, we used the JAR3D program (Zirbel et al., 2015), which scores RNA hairpin and internal loop sequences against motif groups from the RNA three-dimensional motif atlas (Petrov et al., 2013), to search for conserved geometries (other than that of the hammerhead fold) in the sequences of the ribozyme domains of both polarities in the five currently known members of the family *Avsunviroidae* (Di Serio et al., 2018). Interestingly, all ribozyme sequences that were analyzed showed significant matches with the tRNA fold. In its most common configuration, the anticodon hairpin is composed by a 7-nt loop closing a 5-nt stem. The first two nucleotides of the loop are highly conserved as YU (being the CU pair more common than UU). The pre-tRNA intron is spliced-out from nucleotides R and N after the variable anticodon triplet (being A more common than G in the first case, and a preponderance of A in the second) (Fig. 3A, left). The uridine of the conserved sequence YU induces a characteristic U-turn–a rigid sharp turn of the polynucleotide backbone between the U and the first nucleotide of the anticodon–that is necessary to present the anticodon trinucleotide (Robertus et al., 1974). These overall characteristics seem to be conserved in the U-turn loop of the viroid hammerhead ribozymes, located between helices I and II (Fig. 3A, right). This U-turn loop contains the conserved catalytic sequence CUGAYGA (Doudna and Cech, 1995; Pley et al., 1994). This sharp turn in the ribozyme phosphate backbone seems to allow the correct positioning of the three helixes, accommodating the core nucleotides in the appropriate places for the self-cleavage reaction. We reasoned that this turn must have an important role generating the structure that is recognized by tRNA ligase.

**FIG 3.**
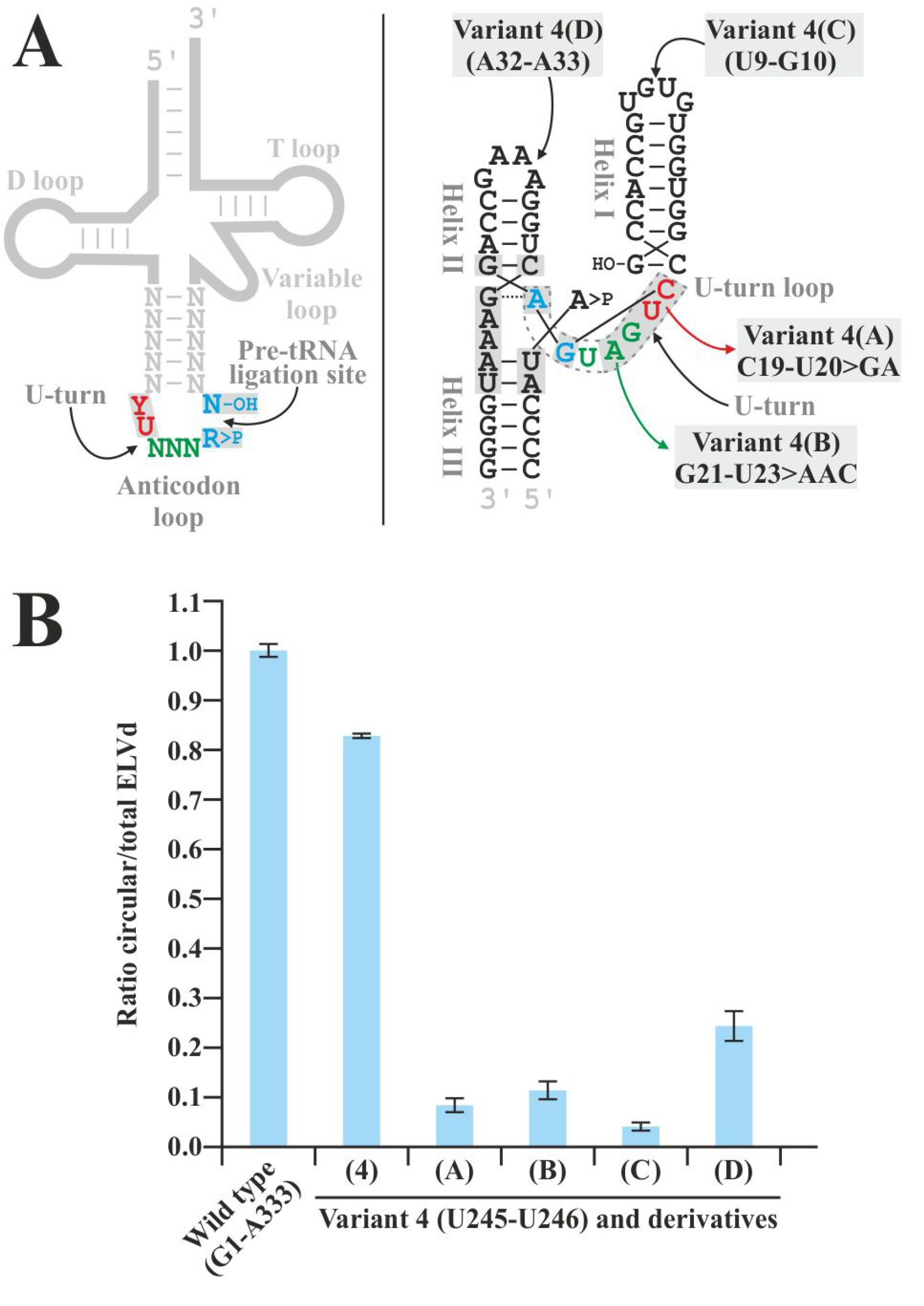
Analysis of the hammerhead ribozyme domain sequences that are relevant during the eggplant tRNA ligase-mediated ligation process. (**A**) Generic cloverleaf structure of tRNAs, focusing the anticodon loop (left) and the ELVd (+) hammerhead ribozyme functional structure during self-cleavage (right). Conserved nucleotides of each RNA are shown in light grey boxes. The analogous nucleotides between both RNAs share colors. ELVd Variants 4A and B, with nucleotide substitution C19-U20>GA and G21-U23>AAC, and 4C and D, opened in different positions in the ribozyme (U9-G10 and A32-A33, respectively), are indicated. (**B**) ELVd variants were expressed in *E. coli* along with the eggplant tRNA ligase. Total RNA was 2D separated and analyzed by northern blot hybridization. Histogram shows the circularization rates (monomeric circular versus monomeric circular plus linear). Error bars represent the standard deviations in three independent *E. coli* clones.

In light of this rational, we designed a set of mutations aimed at testing the mimicry hypothesis, focusing on modifying the set of nucleotides within this U-turn loop that can be relevant in the sharp turn and maintenance of the structure, while being close enough to interact with the cleaved nucleotides. We tried to avoid substantial modifications in the overall secondary structure that the ribozyme acquires during the ligation process. We made these modifications on Variant 4, flanked by two ELVd ribozyme copies. This form is circularized at high levels (Fig. 1C), which makes it possible to analyze the role of the ribozyme without the additive effect of the presence of the domain of the ribozyme of complementary polarity at the natural ligation site. Since modifications in the ribozyme domain have detrimental effects on the correct processing of the precursor RNA, we employed the engineered ribozymes strategy to generate the ligatable termini (Supplementary Dataset S1). We first modified the conserved nucleotides C19-U20 to GA via site-directed mutagenesis (Fig. 3A, right, Variant 4A). These two positions in the hammerhead ribozyme sequence are analogous to the dinucleotide that contains the U-turn and precedes the anticodon in the tRNA. We also modified the conserved nucleotides G21-A22, together with the non-conserved U23, to AAC (Fig. 3A, right, Variant 4B); these three nucleotides are the equivalent of the anticodon triplet. The longer-than-unit ELVd precursors containing these mutations were co-expressed in *E. coli* with the eggplant tRNA ligase; total RNA was separated using 2D-PAGE and analyzed by northern blot hybridization. Interestingly, a drastic reduction in circularization was induced by modifying both sets of nucleotides; the circularization rates fell to approximately 8% and 11% compared to the wild type when the conserved CU or the three GAU nucleotides were modified, respectively (Fig. 3B). Additionally, we tested the functionality of the U-turn in the enzymatic ligation of terminal nucleotides that are located far from it by shifting the ligation point away but keeping it in the ribozyme sequence. The two new opening sites were located within the terminal loops of helices I and II (between positions U9-G10 and A32-A33, respectively) (Fig. 3A, right, Variants 4C and 4D). Northern blot analysis of the total RNA from recombinant bacteria separated by 2D-PAGE showed a reduction to approximately 4 and 24% in the circularization when the ligation site is located in the helix I and II, respectively (Fig. 3B). Altogether, these results support a role for the ribozyme U-turn loop and, more specifically, for some of its conserved residues, not only in ribozyme self-cleavage but also in the circularization process. Considering that these nucleotides are conserved when compared to those present in the tRNA anticodon loop and appear to acquire the same general structure, we posit that a structural mimicry of tRNA is probably occurring to recruit the host tRNA ligase that joins terminal residues located in the vicinity of the internal loop.

## DISCUSSION

In viroid RNA molecules, all of the sequences and structures that are essential to completing replicative cycles are densely packed into small genomes. Due to this tight packing, some motifs likely perform multiple functions and participate in several processes of infection; they are probably recognized by various cellular structures or proteins in the host plant. An example of this functional multiplicity is the loop E motif, which is present in the CCR of the potato spindle tuber viroid. In addition to its canonical role in viroid processing and ligation (Diener, 1986; Gas et al., 2007), loop E has also been associated with the regulation of transcript levels for both, modulate the dynamics of infection (Adkar-Purushothama and Perreault, 2020) and host adaptation (Qi and Ding, 2002; Wassenegger et al., 1996), and with symptom induction (Qi and Ding, 2003). Either loop E submotifs or various transient secondary structures must be responsible for interaction with different host factors (Qi and Ding, 2003). Similarly, in the case of ELVd, while hammerhead ribozymes are involved in the processing of viroid oligomeric transcripts to produce monomeric units, previous research suggests that they may also be involved in the circularization process by adopting a hypothetical transitory structure that favors ligation (Cordero et al., 2018).

To better understand the role of hammerhead ribozyme in tRNA ligase-mediated ELVd circularization, we used an *E. coli*-based experimental system to analyze the ligation of reorganized ELVd monomers with ligatable 5’-hydroxyl and 2’,3’-cyclic phosphodiester terminal groups. Most reorganized forms circularized at substantially lower rates than the genuine linear intermediate (Fig. 1). This result is in agreement with a previous *in vitro* analysis of these same mutants (M.-A. Nohales et al., 2012). Next, we assessed the effect of inserting the ribozyme halves in the different opening sites. The eggplant tRNA ligase-mediated circularization rate significantly increased in all forms when the opening site included the ELVd ribozyme halves (Fig. 1). These results constitute strong evidence for the role of the ribozyme sequences in ELVd ligation.

Next, taking Variant 2 with flanking ELVd hammerhead ribozymes as a case study, we analyzed whether bordering sequences next to the ribozyme domain further favor circularization. Variant 2 shares some properties with the wild-type linear replication intermediate; the opening site is also found next to a loop in the middle of a quasi-double-stranded structure (Fig. 2A). However, Variant 2 is poorly circularized in the absence of terminal ELVd ribozyme sequences (Fig. 1C). The results again showed improved ligation in all cases, including when sequences corresponding to the domain of the - polarity hammerhead ribozyme were added to mimic the situation in the wild-type linear intermediate (Fig. 2). The location of the ligation site in a central position of the viroid molecule in a quasi-rod-like structure (which results from the hybridization of both hammerhead domains) is a common feature in the family *Avsunviroidae* (Giguère et al., 2014). Both strands may exist in a steady state between the compact rod shape and alternative foldings that permit access to the tRNA ligase, possibly facilitated by the sequences surrounding the ribozyme.

After experimental confirmation that the terminal sequences and the domain of the hammerhead ribozyme with - polarity are important for efficient ligation, we searched for an alternative folding that may explain these results using the JAR3D program. Interestingly, homology-based modelling revealed that the internal U-turn loop C19-A25 of the hammerhead ribozyme can fold in a similar fashion to the conserved anticodon loop of tRNAs (Fig. 3A). Mutational analysis of the ELVd flanking ribozymes in Variant 4 supports the importance of these conserved sequences. On one hand, two modifications in the ribozyme’s highly conserved CUGA and GA motifs (along with the non-conserved U residue), drastically reduced circularization to similar amounts in both cases (Fig. 3B). Since this reduction occurs without noticeable variation in the accumulation of linear intermediates with respect to the same variants without mutations in the ribozyme, these results suggest that some of the conserved sequences of the ribozyme catalytic core also play a role in circularization. The decrease in circularization when modifying the equivalent of the anticodon triplet is remarkable, since greater flexibility could be expected given that these nucleotides are variable in the canonical substrates (the tRNAs) of the enzyme. Whether hammerhead ribozymes have adapted to efficiently mimic a particular codon or whether these modified nucleotides are also essential for maintaining the structure recognized by the ligase remains unsolved. On the other hand, maintaining this conserved sequence but moving the ligation site away from it dramatically reduced circularization (Fig. 3B). Based on the predicted secondary structure of the ribozyme (Fig. 3A, right), the *bonafide* ligation site is located in the vicinity of the U-turn loop that shares structural homology with the anticodon loop, while the new opening sites (located in both helices I and II loops) are far apart from this loop. Since the circularization efficiencies between the two mutants were different, we can infer that the ability of the phosphate backbone to restructure and reposition both terminal nucleotides very close to the U-turn loop with greater or lesser ease (helix I and II, respectively), affects the catalytic capacity of the enzyme. Therefore, the proximity of the terminal residues to the loop seems to be an important factor in the ligation process.

tRNA primary transcripts (pre-tRNAs) are extensively modified after transcription. One of these modifications is performed by tRNA splicing endonuclease, which is responsible for releasing a short intron that, in eukaryotes, usually resides in the anticodon loop between nucleotides 37 and 38; it generates two tRNA halves containing 2’,3’-cyclic phosphate and 5’-hydroxyl ends (Yoshihisa, 2014). In plants, both halves are healed and sealed by the multiple activities of the tRNA ligase (Englert and Beier, 2005). Although splicing of tRNAs, either cytoplasmic or organellic, occurs primarily in the cytoplasm, tRNA ligase reportedly also targets other organelles, such as chloroplasts (Englert et al., 2007). This enzyme has been reported to be involved in the repair mechanism for damaged tRNAs, in the circularization of viroids of the family *Avsunviroidae*, and in other non-conventional RNA splicing functions (Englert et al., 2007; Nagashima et al., 2016; M.-A. Nohales et al., 2012). For example, the *Arabidopsis thaliana* tRNA ligase is involved in initiating zygote division, which may be mediated not only by pre-tRNA processing but also by some unconventional splicing reactions (Yang et al., 2017). This same enzyme has been associated with the stress response to unfolded proteins (Mori et al., 2010; Nagashima et al., 2016; Peschek et al., 2015), as it mediates in the splicing of an mRNA encoding a stress-specific transcription factor. Some researchers have speculated that tRNA ligase can recognize certain RNA structures conserved between species (Mori et al., 2010). In addition, thermal denaturation abolishes ligation, although it remains unknown whether this is caused by moving the ligatable termini away or by interrupting secondary structures that the enzyme could recognize (Peschek et al., 2015). Finally, this enzyme is presumed to remain bound to the spliced RNA–as occurs with the exon junction complex– and to interact with the translation machinery (Mori et al., 2010). A similar ribonucleoprotein complex has also been proposed between tRNA ligase and the viroid when co-expressed in *E. coli* (Daròs et al., 2018).

Although it is not known exactly how recognition between the tRNA ligase and the tRNA halves occurs, our work may shed light on both the pre-tRNA processing and the viroid circularization. Several enzymes are known to include domains that interact with specific anticodon loops; these include aminoacyl-tRNA synthases (Rubio Gomez and Ibba, 2020). However, in the case of tRNA ligase, a single enzyme must recognize tRNAs with different anticodons. Therefore, it is expected that this interaction would not be highly restrictive to particular sequences, instead relying on recognizing a common feature, such as that one that apparently can generate the U-turn. This recognition flexibility would have been exploited by non-canonical RNA substrates such as viroids. Similar mimicry relationships may also be used by other RNAs ligated by tRNA ligase; therefore, structural characteristics similar to those described here can be expected in other cellular RNAs.

Viroids adapted to replicating autonomously with minimum sequences, and ribozymes have the capacity to catalyze the backwards self-ligation reaction, possibly via conformational changes (Canny et al., 2007; Nelson et al., 2005). Therefore, it is a paradox that viroid self-ligation capacity remains very inefficient and the circularization is mediated by a host enzymatic activity (Martínez et al., 2009; M.-A. Nohales et al., 2012). Possibly, as viroids became obligate and exclusive parasites of plants, viroid hammerhead ribozymes lost their ability to acquire the adequate conformation for efficient self-ligation, adapting to host ligases by remodelling their variable regions while maintaining the residues strictly required for self-cleavage.

In conclusion, we propose here that viroid hammerhead ribozymes have evolved to mimic the structure of endogenous cellular RNAs that are native substrates for the host tRNA ligase. These ribozymes probably adopt an alternative transitory folding (mimicking that of the tRNA anticodon loop), being able to recruit the enzyme while maintaining its self-cleavage function. In this way, they mediate two steps required in viroid RNA processing: cleavage of multimeric transcripts and circularization of the resultant monomers. The mechanistic model that summarizes the results and observations of this work is shown in Figure 4. Although we have focused on ELVd circularization, what is described here may be a common feature for all members of the family *Avsunviroidae*, considering that the key elements of ribozyme sequence and structure are conserved among species.

**FIG 4.**
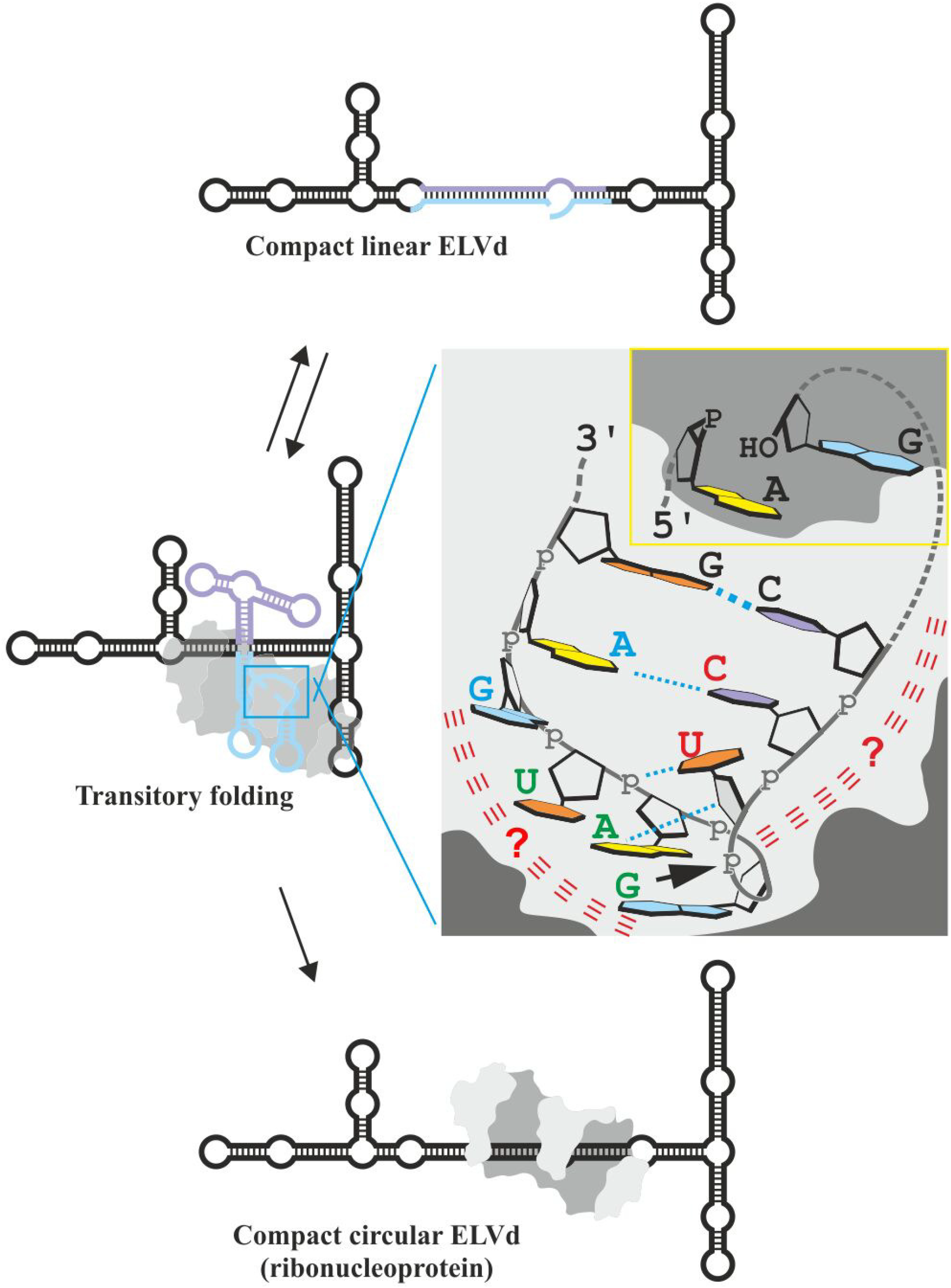
Proposed model of the ELVd circularization mechanism. The natural viroid ligation site is located in a quasi-double-stranded structure formed by the hybridization of + and - hammerhead ribozyme domains in the central region of the molecule. It is likely that this region could form an alternative, less compact structure in which a characteristic folding of both ribozyme domains allows the access of the eggplant tRNA ligase to the terminal nucleotides for the ligation. The model proposes a role for the ribozyme U-turn loop in the enzymatic ligation process. This loop may adopt a fold similar to that of the tRNA anticodon loops that is recognized by the tRNA ligase via unknown interactions (red lines). In the model, the terminal nucleotides, located in the proximity of the catalytic pocket, are correctly positioned in the ligase catalytic center (yellow insert) for its ligation.

## MATERIALS AND METHODS

### Construction of plasmids to express ELVd variants in *E. coli*

Several variants of the ELVd reference sequence (GenBank accession number NC_039241.1) were expressed in *E. coli* from plasmid pLELVd-2 (Daròs et al., 2018). In this plasmid, a longer-than-unit ELVd RNA with + polarity, flanked by two copies of the hammerhead ribozyme (positions C327 to G46), is expressed under the control of the *E. coli* murein lipoprotein promoter and the 5S rRNA (*rrnC*) terminator. This plasmid contains a pUC replication origin and a selection marker that confers ampicillin resistance (Daròs et al., 2018). The mutations in the ELVd sequence contained in this plasmid were created via standard molecular biology techniques. DNA amplifications were performed by polymerase chain reaction (PCR) using the Phusion high-fidelity DNA polymerase (Thermo Scientific). PCR products of the appropriate size were electrophoretically separated in 1% agarose gels and recovered by elution. When required, oligonucleotide primers or PCR products were phosphorylated with T4 polynucleotide kinase (Thermo Scientific). Plasmid assembly was performed by ligation with T4 DNA ligase (Thermo Scientific) or by the Gibson reaction using the NEBuilder HiFi assembly master mix (New England Biolabs). *E. coli* DH5α cells were electroporated with the resulting plasmids; the recombinant clones were selected on plates with lysogeny broth (LB) medium containing 50 μg/ml ampicillin. The plasmids that contained the desired mutations were selected using antibiotic resistance, electrophoretic analysis, and sequencing (3130xl Genetic Analyzer, Life Technologies).

### Co-expression of the ELVd variants and eggplant tRNA ligase in *E. coli*

To study circularization of the different ELVd variants, *E. coli* DH5α cells were co-electroporated with pLELVd-2 derivatives (see above) and p15LtRnlSm (Daròs et al., 2018), which encodes the chloroplastic isoform of the eggplant tRNA ligase (GenBank accession no. JX0225157). The expression of this protein is also under the control of the *E. coli* murein lipoprotein promoter and the *rrnC* terminator; the plasmid contains a p15A replication origin and a chloramphenicol selection marker (Daròs et al., 2018). Recombinant colonies were selected for in plates with 50 μg/ml ampicillin and 34 μg/ml chloramphenicol. Isolated colonies were inoculated into 50 ml tubes with 5 ml of LB medium containing both antibiotics and were grown for 24 h at 37°C with vigorous shaking (225 revolutions per min).

### RNA extraction

Aliquots (2 ml) of the cultures were taken at the indicated times. Cells were pelleted by centrifugation at 13,000 rpm for 2 min and resuspended in 50 μl of TE buffer (10 mM Tris-HCl, pH 8.0, 1 mM EDTA). One volume of a 1:1 (v/v) mixture of phenol (saturated with water and equilibrated to pH 8.0 with Tris-HCl, pH 8.0) and chloroform was added; the cells were lysed by vigorous vortexing. After centrifugation at 13,000 rpm for 5 min, the aqueous phases with the total bacterial RNA were recovered. They were subjected directly to analysis or were stored at −20°C.

### RNA analysis

Total *E. coli* RNA was separated by 2D-PAGE (Daròs, 2022; Ortolá et al., 2021). Aliquots (20 μl) of the aqueous phases were mixed with 1 volume of loading buffer (98% formamide, 10 mM Tris-HCl, pH 8.0, 1 mM EDTA, 0.0025% bromophenol blue, and 0.0025% xylene cyanol); they were incubated for 1.5 min at 95°C and snap-cooled on ice. RNA was separated under denaturing conditions in 5% polyacrylamide gels (37.5:1 acrylamide:N,N-methylenebisacrylamide) in TBE buffer (89 mM Tris, 89 mM boric acid, and 2 mM EDTA) with 8 M urea. The electrophoresis was carried out at 200 V for 2 h; the gels were then stained with agitation in 200 ml of 1 μg/ml ethidium bromide for 15 min. After washing three times with water, fluorescence in the gel was recorded under UV light (UVIdoc-HD2/20MX, UVITEC). The lanes of interest were then cut and placed transversely on top of a series of denaturing (8 M urea) 5% polyacrylamide gels casted as described above, but in 0.25X TBE buffer. These electrophoreses were run at 350 V (maximum 25 mA) for 2.5 h, and the gels were stained and recorded as described above.

After 2D-PAGE separation, RNA was electroblotted to positively charged nylon membranes (Nytran SPC, Whatman) and crosslinked with 1.2 J/cm^2^ UV light. Membranes were then subjected to hybridization overnight at 70°C with hybridization buffer (50% formamide, 0.1% Ficoll, 0.1% polyvinylpyrrolidone, 100 ng/ml salmon sperm DNA, 1% sodium dodecyl sulfate –SDS–, 0.75 M NaCl, 75 mM sodium citrate, pH 7.0) containing approximately 1 million counts per minute of a ^32^P-labeled monomeric ELVd RNA probe with - polarity. The ELVd probe was obtained via *in vitro* transcription of a linearized plasmid for 1 h at 37°C with 20 U of T3 bacteriophage RNA polymerase (Roche) in 40 mM Tris-HCl, pH 8.0, 6 mM MgCl_2_, 20 mM DTT, 2 mM spermidine, 0.5 mM each of ATP, CTP and GTP, 50 μCi of [α-^32^P]UTP (800 Ci/mmol), 20 U RNase inhibitor (RiboLock, Thermo Scientific), and 0.1 U yeast inorganic pyrophosphatase (Thermo Scientific). After transcription, the linearized plasmid was digested with 20 U DNase I (Thermo Scientific) for 10 min at 37°C. The radioactive probe was purified chromatographically with a Sephadex G-50 column (mini Quick Spin DNA Columns, Roche). After hybridization, the membranes were washed three times for 10 min at room temperature with 2X SSC (1X SSC is 150 mM NaCl, 15 mM sodium citrate, pH 7.0), 0.1% SDS; they were washed once for 15 min at 55°C with 0.1X SSC, 0.1% SDS. Hybridization signals were recorded in an imaging plate (BAS-MP, FujiFilm) and quantified using an image analyzer (Amersham Typhoon, GE Healthcare).

### Computational analysis

The minimum free energy conformation of the various monomeric linear ELVd variants was predicted using the CentroidFold algorithm (Sato et al., 2009). Homology between RNA loops was analyzed using the JAR3D algorithm (Zirbel et al., 2015).

## ACKNOWLEDGEMENTS

This research was supported by grants BIO2017-91865-EXP and PID2020-114691RB-I00 from Ministerio de Ciencia e Innovación (Spain) through the Agencia Estatal de Investigación, co-financed by European Regional Development Fund. B.O. was the recipient of a pre-doctoral contract (PAID-01-17) from Universitat Politècnica de València.

B.O and J.A.D. designed the research. B.O. performed the experiments. B.O and J.A.D analyzed the results and wrote the article.

